# Adaptations during maturation in an identified honeybee interneuron responsive to waggle dance vibration signals

**DOI:** 10.1101/469502

**Authors:** Ajayrama Kumaraswamy, Hiroyuki Ai, Kazuki Kai, Hidetoshi Ikeno, Thomas Wachtler

## Abstract

Honeybees are social insects, and individual bees take on different social roles as they mature, performing a multitude of tasks that involve multi-modal sensory integration. Several activities vital for foraging, like flight and waggle dance communication, involve sensing air vibrations using antennae. We investigated changes in the identified vibration-sensitive interneuron DL-Int-1 in the honeybee *Apis mellifera* during maturation by comparing properties of neurons from newly emerged and forager honeybees. Comparison of morphological reconstructions of the neurons revealed minor changes in gross dendritic features and consistent, region dependent and spatially localized changes in dendritic density. Comparison of electrophysiological properties showed an increase in the firing rate differences between stimulus and non-stimulus periods in foragers compared to newly emerged adult bees. The observed differences in neurons of foragers as compared to newly emerged adult honeybees indicate refined connectivity, improved signal propagation, and enhancement of response features important for the network processing of air vibration signals relevant for the waggle-dance communication of honeybees.

## Introduction

Perception of vibrations and sounds is very important for social insects (Hunt and Richard 2013) and among them, honeybees are unique in that they use air-borne vibrations in addition to substrate-borne vibrations for communication (Kirchner 1997). Among several intra-hive communication behaviors linked to air-borne vibration sensing (Barth et al. 2005; Hunt and Richard 2013; Nieh 2010), the waggle dance behavior, by which honeybees communicate the location and profitability of food sources, has been extensively studied (von Frisch 1965; von Frisch 1993; Kirchner 1993; Brockmann and Robinson 2007; Hrncir et al. 2011; Couvillon 2012). However, the neural mechanisms underlying the processing and decoding of the waggle dance vibration signals have so far not been uncovered.

Honeybees detect air vibrations using Johnston’s organ (JO) located in the pedicel of their antennae (Figure 1a; Dreller and Kirchner 1993). Sensory afferents of the JO project into the dorsal lobe (DL), dorsal subesophageal ganglion (dSEG) and medial posterior protocerebral lobe (mPPL) of the honeybee brain (Figure 1a; Ai et al. 2007).

**Figure 1.**
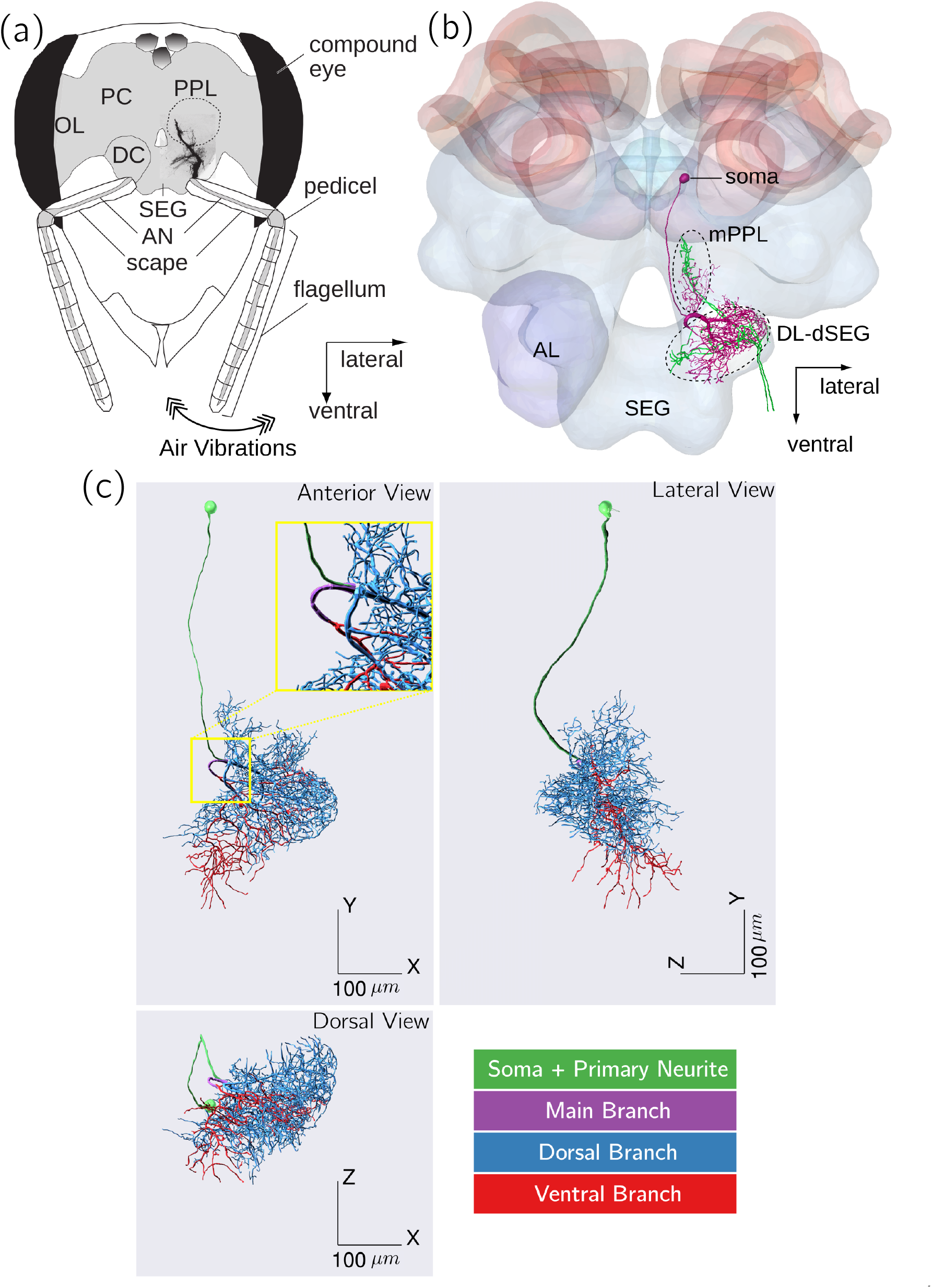
Vibration Sensing, primary auditory center and DL-Int-1 interneuron in the honeybee. **(a)** Airborne vibrations picked up by the flagellum are transduced by sensory neurons of the Johnston’s organ (JO) in the pedicel and transmitted to the primary auditory center of the honeybee brain, which consists of medial PPL, DL and dorsal SEG. Modified version of Figure 1 of Ai et al. (2007). **(b)** Projection patterns of sensory afferents (green) and DL-Int-1 in the primary auditory center of the honeybee brain. DL-Int-1 has dendrites running close to sensory afferents in the DL. Modified version of Figure 5 of Ai (2013). **(c)** Morphology of DL-Int-1 visualized using three 2D projections. We divided DL-Int-1 morphology into four subregions for analyses. Inset: A magnified version of the region around the Main Branch. *OL: optic lobe, PC: protocerebrum, DC: deutocerebrum, PPL: posterior protocerebral lobe, SEG: subesophageal ganglion, AN: antennal nerve, AL: antennal lobe, mPPL: medial PPL, dSEG: dorsal SEG*

More than ten groups of interneurons belonging to three categories have been identified and characterized in these regions (Ai et al. 2009; Ai 2010; Ai 2013; Ai et al. 2017)) and have been shown to respond to air-borne vibration pulses similar to those produced during the waggle dance (Ai et al. 2017). In particular, a group of GABAergic interneurons called DL-Int-1 has been studied intensively and has been characterized in detail (Ai et al. 2009; Ai 2013; Ai et al. 2017).

DL-Int-1 somata are located in the rind of the protocerebrum (PC) and have single neurites branching and projecting to the DL, the dSEG and the mPPL, where they further branch into dense arborizations that run close to JO afferents (Figure 1b; Ai et al. 2009). DL-Int-1 are spontaneously active and respond to vibration stimuli applied to the ipsilateral antenna. Their responses to vibration stimuli are characterized by on-phasic excitation to stimulus onset, tonic inhibition during continuous stimulation, and rebound spiking after vibration offset (Ai et al. 2009). DL-Int-1 neurons are thought to play a central role in encoding the duration of the waggle phase (Ai et al. 2017), which indicates the distance of the advertised food source from the hive (von Frisch 1993).

As they mature, adult honeybees engage in four primary social roles—cleaners, nursers, food storers and foragers—and perform different tasks in different roles (Seeley 1996). Several studies have investigated the neural basis of such behavioral versatility by studying structural changes in the honeybee brain with age and social role, mainly focusing on the mushroom body (Meinertzhagen 2010; Groh et al. 2006; Groh et al. 2012). Although most developmental changes in the honeybee brain occur during pupal and larval stages (Devaud and Masson 1999), considerable age-dependent and experience-dependent anatomical changes have been described at the level of subregions in the adult honeybee antennal lobe (Winnington et al. 1996; Sigg et al. 1997; Morgan et al. 1998; Brown et al. 2004; Andrione et al. 2017; Arenas et al. 2013) and the mushroom body (Withers et al. 1993; Withers et al. 1995; Durst et al. 1994; Fahrbach et al. 1998; Wolschin et al. 2009), as well as at the level of single mushroom body neurons (Farris et al. 2001). In addition, electrophysiological properties of honeybee neurons also mature with age and experience in the antennal lobe (Wang et al. 2005) and in the mushroom body (Kiya et al. 2007). Adaptations in neural processing could be especially crucial during the transition to foraging, since, in comparison to in-hive activities, foraging entails several new and complex behaviors such as attending to waggle dancers, sensing the waggle dance vibration signals and decoding target location, using such information on foraging trips, and advertising newly found locations to hivemates through the waggle dance. It is unclear, however, to what extent neurons in central circuits processing waggle dance vibration signals show such adaptation. We therefore investigated morphological and electrophysiological changes of neurons in the primary auditory center of the honeybee, focusing on DL-Int-1 neurons. To identify maturation-related adaptations in DL-Int-1, we analyzed and compared reconstructed morphologies and electrophysiological properties of neurons from newly emerged adult bees and foragers bees.

## Materials and Methods

### Honeybees

Honeybees (*Apis mellifera*) reared at Fukuoka University between 2012 and 2014 were used in this study. Experiments were conducted on more than 300 bees for investigating neurons in the primary auditory center of the honeybee brain. Collected data included electrophysiological recordings and laser scanning microscopy images, which were stored in a database and classified into multiple neuron groups based on electrophysiological and morphological characteristics (see Table 1, Ai et al. 2017). In the current study, we used DL-Int-1 data from the database belonging to honeybees of two stages of maturation:

- **Newly emerged adults** (age 1-3 days): female honeybees shortly after emerging from their cell in the hive. Before the experiments, these bees were kept in isolated cages containing sugar solution and pollen.
- **Foragers** (older than 10 days): female honeybees returning from foraging with pollen on their hindlegs.

**Table 1.** Scalar morphometric measures showing significant differences. Summary statistics of five scalar morphometric and topological parameters that show significant differences for at least one morphological subregion. P-values were calculated using Welch’s unequal variances t-test and a cutoff of 5% was used. Summary statistics for all nineteen scalar measures and for all four subregions are provided in Table S1, Table S2, Table S3 and Table S4 of Supplementary Data. Morphologies show significant differences for a few scalar measures, with width, total dendritic volume and maximum centrigual order showing large changes.

| Measure | Morphological Subregion | Newly Emerged (NE, n=6) | Forager (F, n=6) | % change in mean value $\{(F-NE)/NE\}$ | p-Value |
| --- | --- | --- | --- | --- | --- |
| Width (along X) ($\mu m$) | main branch | $20.5 \pm 5.27$ | $39.1 \pm 13.6$ | +90.6% | 0.026 |
| Width (along X) ($\mu m$) | dorsal branch | $215 \pm 30.0$ | $259 \pm 30.6$ | +20.5% | 0.045 |
| Total dendritic volume ($\times 10^3(\mu m)^3$) | main branch | $0.431 \pm 0.0202$ | $0.763 \pm 0.0220$ | +77.0% | 0.033 |
| Average partition asymmetry | dorsal branch | $0.619 \pm 0.013$ | $0.583 \pm 0.031$ | -5.82% | 0.046 |
| Average partition asymmetry | whole arborization | $0.616 \pm 0.0180$ | $0.584 \pm 0.0252$ | -5.2% | 0.045 |
| Maximum centrifugal order | ventral branch | $29.7 \pm 10.2$ | $15.8 \pm 2.34$ | -51.6% | 0.028 |
| Hausdorff fractal dimension | ventral branch | $1.24 \pm 0.0706$ | $1.15 \pm 0.0339$ | -7.26% | 0.035 |

### Experimental Procedure

The experimental procedure for generating LSM image stacks and electrophysiological response traces from honeybee vibration-sensitive interneurons has been presented in detail in Ai et al. (2017) and we describe it here briefly. After immobilization and head fixing using bee’s wax, the frontal surface of the honeybee brain was exposed by cutting away a small rectangular window between the compound eyes. Borosilicate glass electrodes filled at the tip with a dye were inserted into the DL to record from individual neurons. Three dyes were used — Lucifer Yellow CH dilithium salt (catalog number L0259, Sigma-Aldrich), Dextran Tetramethylrhodamine solution (3000 molecular weight, anionic, lysine fixable; catalog number D3308, Thermo Fisher Scientific) and Alexa Fluor 647 hydrazide (catalog number A20502, Thermo Fisher Scientific). With the electrode stably inserted into an vibration-sensitive interneuron, sinusoidal vibration stimuli at 265 Hz with 1 s duration were applied to the right antenna and responses were recorded intracellularly. Electrical signals were amplified using an amplifier (MEZ8301, Nihon Kohden) and recorded using Spike2 (Cambridge Electronic Design; `RRID: SCR_000903`). After recording electrical activity, a hyperpolarizing current (2–5 nA for 2–10 min) was applied to inject the dye into the neuron. Thereafter, the brains were dissected out, fixed in 4% paraformaldehyde for 4 h at room temperature, then rinsed in phosphate buffer solution, dehydrated, and cleared in methyl salicylate for subsequent observation and imaging.

The cleared specimens containing intracellularly stained neurons were viewed from the posterior side of the brain under a confocal laser-scanning microscope (LSM 510, Carl Zeiss) with a Zeiss Plan-Apochromat 25x/numerical aperture 0.8 oil lens objective (working distance, 0.57 mm). Image stacks of the DL-dSEG and medPPL were taken at a resolution of 0.36 µm on the imaging plane using 1.0 µm deep optical sections and stitched together digitally to obtain image stacks of complete neurons.

### Morphological Subregions of DL-Int-1

To refer to specific subregions of the DL-Int-1 morphology, we adopt the following definitions (Figure 1c, see also Ai et al. (2017)):

- **soma and primary neurite (SPN):** consists of the soma and its primary neurite until bifurcation
- **main branch (MB):** consists of two daughter branches of the primary neurite until they bifurcate.
- **dorsal branch (DB):** consists of the remaining dendritic arborization originating from the dorsal end of MB.
- **ventral branch (VB):** consists of the remaining dendritic arborization originating from the ventral end of MB.
- **whole arborization (WA):** consists of MB, DB and VB.

### Reconstruction of morphologies

The reconstruction procedure has been described in detail in Ikeno et al. (2018). Briefly, LSM image stacks with single dye-filled neurons were deconvolved to reduce image blurring and noise. Regions of each image stack containing the dendritic subtrees emerging from the dorsal and ventral daughter branches of SPN (Figure 6E of Ikeno et al. 2018) were identified based on continuity of branching structure and dendritic thickness and were converted into custom image masks. Applying these masks, two image stacks were created that separately contained the the identified dorsal and ventral subtrees of SPN. Morphologies of these subtrees were reconstructed from their image stacks by segmentation, pruning and smoothing using the SIGEN software (Minemoto et al. 2009, `RRID: SCR_016284`). The morphology of SPN was reconstructed separately and combined with the reconstructed dendrites of the dorsal and ventral subtrees to obtain the reconstruction of a complete DL-Int-1 neuron.

After reconstruction, the data of each DL-Int-1 morphology were manually separated into SPN and WA and stored in separate SWC files (Cannon et al. 1998). SPN was not further analyzed. The data of WA were manually separated into MB, DB and VB based on the first branching points on the two daughter branches of the primary neurite and stored in separate SWC files for morphometric analyses.

### Analysis of morphology

#### Scalar Measures

Morphologies of DL-Int-1 neurons from newly emerged adult and forager honeybees were compared using nineteen widely used metric and topological measures (Uylings and van Pelt 2002; Scorcioni et al. 2008; Peng et al. 2014). These measures were calculated using a modified version of the BT-MORPH software (Torben-Nielsen 2014, `RRID: SCR_003566`, modified code: https://github.com/wachtlerlab/btmorph_v2, version 2.2.1), Vaa3d (Peng et al. 2014, `RRID: SCR_002609`, version 3.447) and pyVaa3d (https://github.com/ajkswamy/pyVaa3d, version 0.4). For each measure, Welch’s unequal variances t-test (Ruxton 2006; McDonald 2014) was used to calculate the significance of difference in mean values of newly emerged adult and forager morphologies with a cutoff of 5%.

#### Spatial Registration

Preliminary visual comparisons indicated that DL-Int-1 morphologies had differences in translation, rotation and scaling that could have resulted from structural differences between honeybee brains. Therefore, we co-registered all DL-Int-1 morphologies to a common frame of reference using the Reg-MaxS-N software (Kumaraswamy et al. 2018, software version at `doi:10.12751/g-node.feee47`, `RRID: SCR_016257`) by removing translation, rotation and scale differences at spatial scales of 160, 80, 40 and 20 µm. Morphologies from newly emerged adult and forager bees were co-registered in two steps. First, newly emerged adult and forager morphologies were co-registered separately using Reg-MaxS-N (Kumaraswamy et al. 2018). Then, the two resulting groups of morphologies were brought to the same frame of reference by co-registering the unions of the points of all morphologies in a group using Reg-MaxS (Kumaraswamy et al. 2018).

To control for parameter choice during spatial registration, the procedure above was repeated using multiple parameter sets. Newly emerged adult and forager morphologies were each co-registered separately using three initial references to generate three sets of registered morphologies each. Picking one set each for the two maturation levels in all possible ways, nine sets of all twelve morphologies were created, which were in turn registered together. All other parameters remained the same for the nine sets (see Table S5 of Supplementary Data for all parameters).

### Comparison using spherical shells

The radial distribution of dendritic length was compared between the two maturation levels by dividing the space containing the morphologies into spherical shells of thickness 20 µm, similar to Sholl Analysis (Sholl 1953; Uylings and van Pelt 2002; Langhammer et al. 2010; Garcia-Segura and Perez-Marquez 2014), which has been shown to be effective in analyzing morphologies (Cuntz et al. 2008; Luebke et al. 2013; O’Neill et al. 2015). As a natural extension of Sholl analysis, we used dendritic length, which is a basic measure of dendritic arborization, to investigate changes during maturation. For every shell, we calculated the percentage of dendritic length (PDL) of a morphology contained in the shell. Using two-way ANOVA (Fujikoshi 1993; McDonald 2014), we tested whether, in each shell, PDL (1) was significantly different between newly emerged adults and foragers, (2) was not significantly affected by registration parameters and (3) showed no significant dependence between the effects caused by maturation and registration parameters.

The above tests used a cut-off level of significance of 5% after Bonferroni correction (Bland and Altman 1995; McDonald 2014). This analysis was not applied to MB morphologies since most of them had no branching points and comprised of single stretches of dendrites spanning less than 50 µm.

### Comparison using 3D voxels

The morphologies were compared at an even finer spatial scale, considering 3D voxels of size 20 µm. We calculated the sum of the dendritic length contained in each voxel, which we refer to as local dendritic length (LDL). Note that, since all the voxels used had the same volume, changes in LDL are proportional to changes in average dendritic density per voxel. The same criteria as in the previous analysis were used for identifying the voxels for which LDL changed significantly during maturation, independent of registration parameters. To quantify and visualize the changes in dendritic density, we calculated for each voxel the normalized change in LDL (ΔLDL_*norm*_):

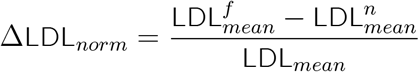

where, for a given voxel, 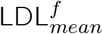 is the average LDL for forager morphologies across registration parameters and honeybee samples, 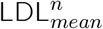 is the average LDL for newly emerged adult morphologies across registration parameters and honeybee samples; and LDL_*mean*_ is the average LDL across all maturation levels, registration parameters and honeybee samples.

Analysis of morphologies using concentric spherical shells and 3D voxels was done using custom Python (`RRID: SCR_008394`) scripts.

### Analysis of electrophysiology

The physiological response of DL-Int-1 to continuous vibration stimuli applied to the antenna consisted of on-phasic excitation followed by a tonic inhibition and offset rebound (Ai et al. 2009). We defined four time periods for analyzing the electrophysiological activity of DL-Int-1 (Figure 4b):

- **Spontaneous Activity:** 3 s period preceding stimulus onset
- **On-phasic Response:** First 75 ms after stimulus onset
- **Inhibitory Response:** From the end of On-phasic Response until stimulus offset
- **Rebound Response:** A 75 ms period after a delay of 25 ms from stimulus offset

Raw data of electrophysiological recordings were read from the Spike2 files using NEO version 0.5 (Garcia et al. 2014, `RRID: SCR_000634`), stored using the NIX format version 1.4.5 (Stoewer et al. 2014, `RRID: SCR_016196`) and analyzed using custom Python scripts. Trials were time-aligned to stimulus onset and time-resolved estimates of average firing rates were generated using adaptive kernel density estimation (Shimazaki and Shinomoto 2010, Implementation: https://github.com/cooperlab/AdaptiveKDE). The distribution of spike train features such as spike rates and spike times were visualized using the “violinplot” function of the Python package seaborn (Waskom et al. 2018). This function uses kernel density estimation to estimate continuous distributions using Gaussian Kernels and Scott’s formula for bandwidth calculation (Härdle et al. 2004, p73). Welch’s unequal variance t-test was used for testing the significance of differences in response features.

We quantified the strength of inhibition relative to spontaneous activity using Relative Inhibition, which was defined as:

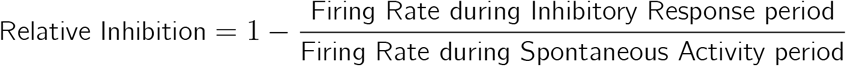

### Computational environment, code and data availability

The analysis was carried out on a desktop computer with an 8-core Intel i7 Processor, 16GB of RAM running Ubuntu 1604. The data used for this study are available online at https://web.gin.g-node.org/The_Ginjang_Project/DL-Int-1_ageBasedChanges_data. The code/software used in this study is available online at https://github.com/wachtlerlab/GJEphys and https://github.com/wachtlerlab/GJMorph.

## Results

### Data Collection

Sharp electrodes were inserted into DL-Int-1 neurons in the honeybee brain to record electrophysiological activity as well as to inject dye for imaging neuron morphology. The success rate of hitting a DL-Int-1 neuron in these experiments was low since honeybee brains were not transparent enough for visually targeted electrode insertion. Furthermore, maintaining the electrode within the neuron long enough to obtain sufficient electrophysiological data was difficult especially for newly emerged adult honeybees, as their brains were soft and infirm. Our data of DL-Int-1 neurons from newly emerged adults were therefore limited to six samples with sufficient data for analysis. For the comparative analysis, we chose six forager samples from our database matching the response pattern of the neurons from newly emerged adults.

### Morphological Adaptations

The four subregions of DL-Int-1 morphology – the whole arborization (WA), main branch (MB), dorsal branch (DB) and ventral branch (VB) (see Methods→Reconstruction of morphologies) were compared separately to investigate changes during maturation.

#### Analysis 1: Scalar Morphometrics

We first compared the morphologies using whole-cell scalar measures, which detect net overall changes in morphological subregions as they combine data from all dendrites. Table 1 lists the measures that showed significant differences between the morphologies of newly emerged adult and forager DL-Int-1 neurons for at least one subregion (see Table S1,Table S2, Table S3 and Table S4 of Supplementary Data for summary statistics of WA, MB, DB and VB respectively). The MB showed significant increases in extent along the X axis (width, 90.7%) and in dendritic volume summed over all dendrites (total dendritic volume, 77.0%). The DB showed a significant increase in its extent along the X axis (width, 20.5%) and a significant decrease in average partition asymmetry (5.8%, Uylings and van Pelt 2002), which quantifies the imbalance between the two subtrees of a branching point. The WA also showed a significant reduction in average partition asymmetry (5.2%). The VB showed significant reductions in maximum centrifugal order (51.6%), which is the maximum number of branch points encountered in a path from root to tip, considering all such paths in a morphology, as well as Hausdorff fractal dimension (7.8%, Smith et al. 1996) which is a measure of the complexity of dendritic arborization.

Thus, significant differences between the neuron morphology of newly emerged adult and forager DL-Int-1 were found in few morphometric parameters. However, these changes were neither consistent across morphological subregions nor highly significant (p-Values between 1% and 5%), indicating that there were no major changes in gross features or branching patterns of DL-Int-1 during maturation at the level of the whole neuron. Nonetheless, they strongly indicated localized changes in dendritic arborization and hence we investigated the morphologies at finer spatial scales, using concentric spherical shells.

#### Analysis 2: Radial distribution of dendritic length

Before detailed spatial analysis, DL-Int-1 morphologies of newly emerged adults and foragers were co-registered to a common frame of reference (see Methods→Analysis of morphology→Spatial Registration and Analysis) to establish spatial correspondence. Figure 2 compares the radial distributions of PDL between newly emerged adults and foragers for WA, DB and VB and highlights those spherical shells for which PDL changed significantly during maturation independent of registration parameters (see Methods→Analysis of morphology→Spatial Registration and Analysis). WA and DB showed reductions in PDL for the shell at 110 µm and increases for shells at 170 and 230 µm during maturation. VB showed reductions at 30–130 µm and increases at 210–270 µm. Thus, there was a consistent reduction of PDL for proximal regions and an increase in distal regions of the morphology.

**Figure 2.**
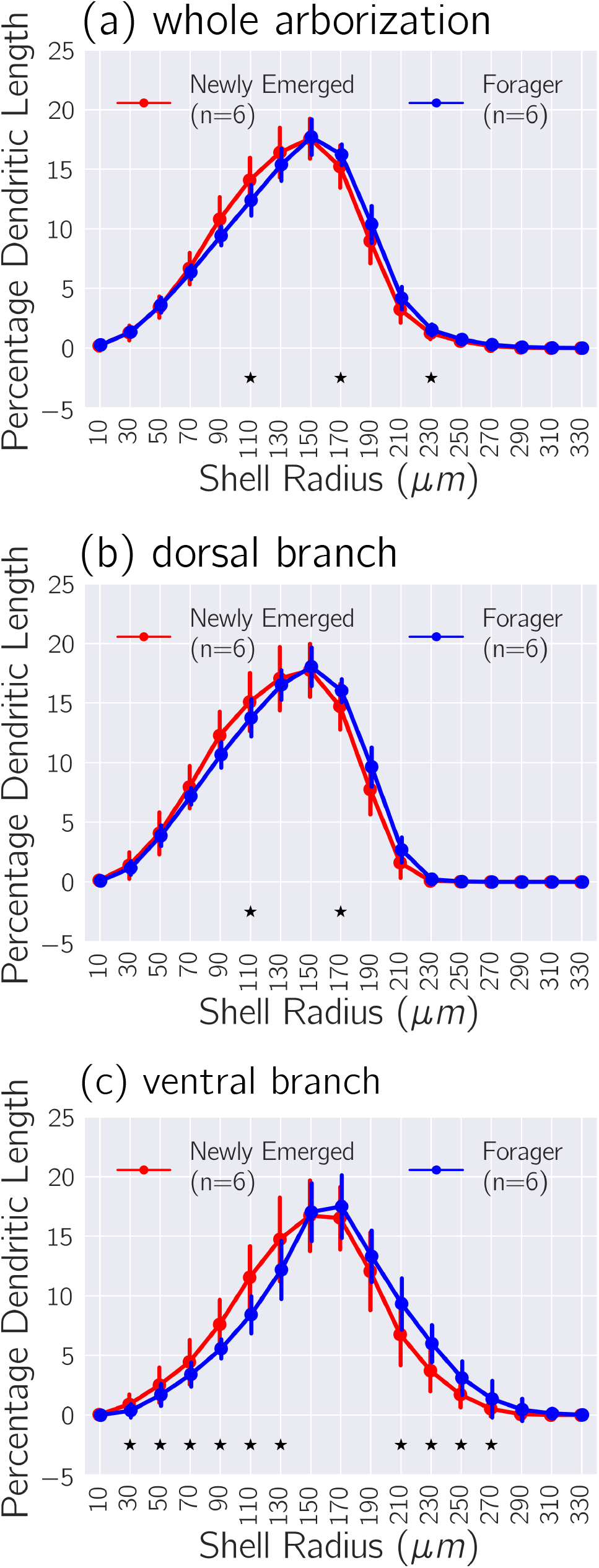
Changes in radial distribution of dendritic density. Comparing the distribution of PDL over concentric spherical shells of thickness 20*µm* for WA, DB and VB in **(a)**, **(b)** and **(c)** respectively. The distributions were calculated by pooling across morphologies generated using different registration parameters. Solid circles indicate mean value and error bars indicate standard deviations. Stars mark the shells for which two-way ANOVA indicated that maturation had a significant effect on PDL, while registration parameters had no such significant effect and the effects of the two factors were not significantly dependent. These comparisons indicate a redistribution of dendritic length during maturation, with pruning in proximal parts and outgrowth in distal parts of DL-Int-1 morphologies.

A pattern of proximal reduction and distal increase of dendritic density would also arise if there was an overall increase in size between the morphologies of newly emerged adults and foragers. However, an overall size difference is unlikely due to our registration method, which includes scaling to achieve a match of the overall volumes occupied by the morphologies, Nevertheless, to exclude that this finding was due to a residual scaling difference, we repeated the co-registration of the morphologies with artificially scaled-up versions of the newly emerged adult morphologies. We scaled up newly emerged adult morphologies by 10 or 15% before bringing all morphologies to the same frame of reference, i.e., between step one and two of the co-registration procedure (see Methods→Analysis of morphology→Spatial Registration and Analysis). That is, the registration processes started with morphologies of newly emerged adult and foragers that had similar spatial spans (in the case of 10% up-scaling), or with morphologies of newly emerged spanning a larger region of space (in the case of 15% up-scaling). For both cases, the maturation related differences in PDL described above remained after registration, in spite of the scaling applied before registration (data not shown). This confirmed that scaling differences did not cause the observed changes in the spatial distributions of dendritic length during maturation, namely, reductions in proximal regions and increases in distal regions.

#### Analysis 3: Local dendritic length

To investigate whether the observed changes in dendritic length distributions were localized in certain subregions of the dendritic arborization, we compared the morphologies at the scale of voxels of size 20 µm using LDL (see Methods→Analysis of morphology→Spatial Registration and Analysis). Figure 3 visualizes the magnitude and spatial distribution of normalized change in LDL (see Methods→Analysis of morphology→Spatial Registration and Analysis) for voxels showing significant changes in the WA, DB and VB using a colormap. Consistent with results from the previous analysis, some proximal regions of the morphologies showed reductions in LDL while some distal regions showed increases. Increases were particularly pronounced in the arborization of the DB in the DL, which showed two main regions (yellow arrows in Figure 3b) where LDL increased during maturation by up to 119% of the average, while decreases were seen in several smaller regions of the DB where LDL decreased by up to 99.2% of the average. In contrast, the VB primarily showed reductions in proximal regions, which were as much as 115% of the average in certain regions, while a few isolated regions showed increases, with a maximum increase of 74.6% of the average (see Figure S1 of Supplementary data for distributions of normalized change in LDL for WA, DB and VB).

**Figure 3.**
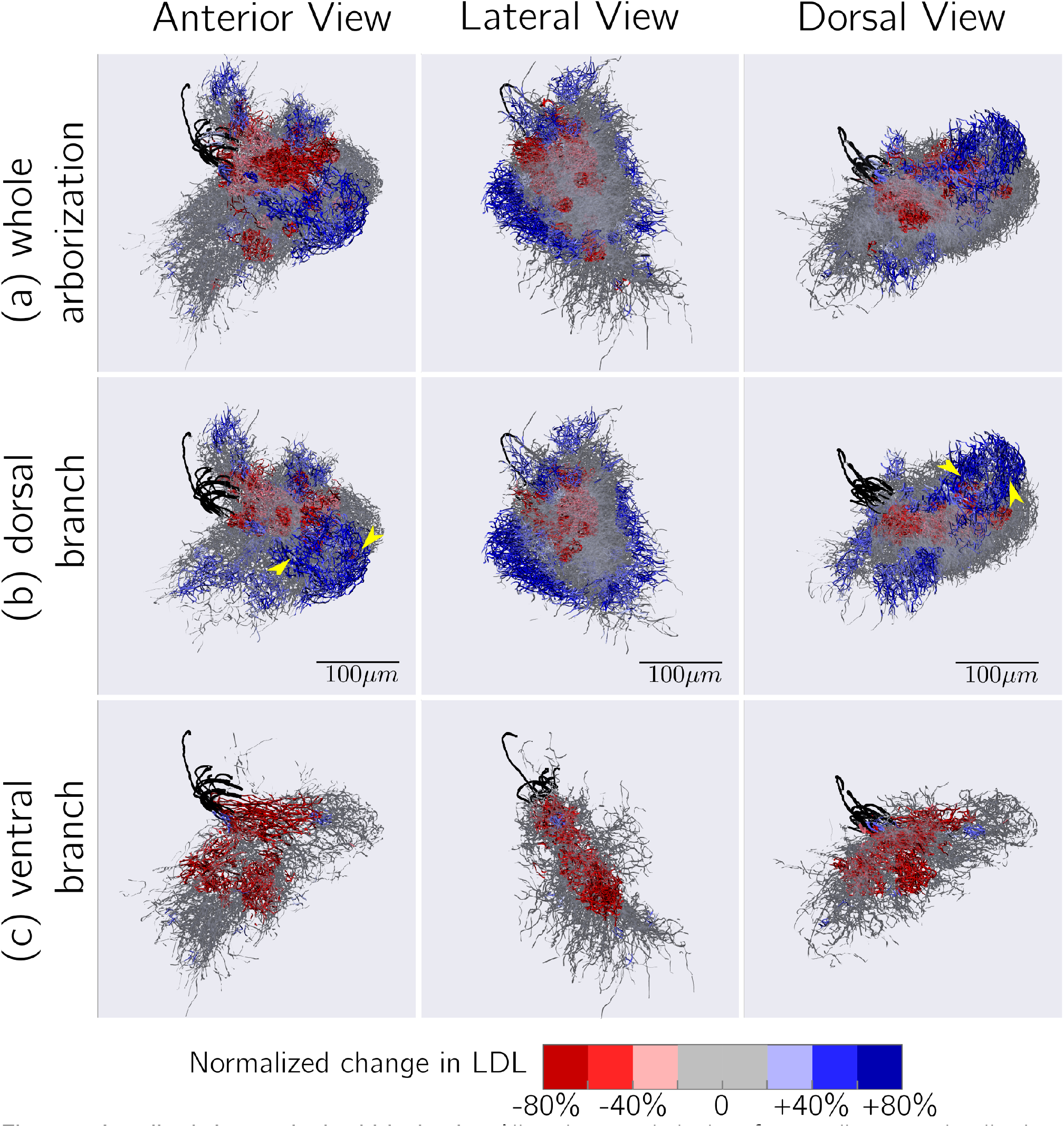
Localized changes in dendritic density. All twelve morphologies after co-alignment visualized together, highlighting regions that show significant changes in LDL between newly emerged adults and foragers. **(a)** whole arborization; **(b)** dorsal branch; **(c)** ventral branch. A voxel was highlighted if two-way ANOVA indicated that maturation had a significant effect on LDL, while registration parameters had no such significant effect and the effects of the two factors were not significantly dependent. The dendrites were colored with normalized change in LDL (see Methods→Analysis of morphology→Spatial Registration and Analysis for definition, see Figure S1 of Supplementary Data for distributions). The Main Branches are colored in black. Yellow arrows in (b) indicate two regions of the arborization of the DB in the DL that show pronounced increases in LDL.

### Electrophysiological Adaptations

Comparison of time-resolved firing rate estimates of the responses of DL-Int-1 neurons (Figure 5a) indicated increased spontaneous activity and a remarkable increase in firing rate just after stimulus offset in foragers as compared to newly emerged adults. These observations were quantified by comparing the firing rates during the four activity periods (Figure 4b) as well as the spike timing during On-phasic response (Figure 4c). Figure 5b summarizes the comparison of firing rates for the four periods. Average spontaneous firing rate showed a significant increase of 39.6%. Average firing rates during on-phasic and inhibitory response periods did not show significant changes, but average firing rate during rebound response nearly doubled, increasing by 94.75%. Thus, DL-Int-1 responses showed stronger spontaneous activity and rebound response.

**Figure 4.**
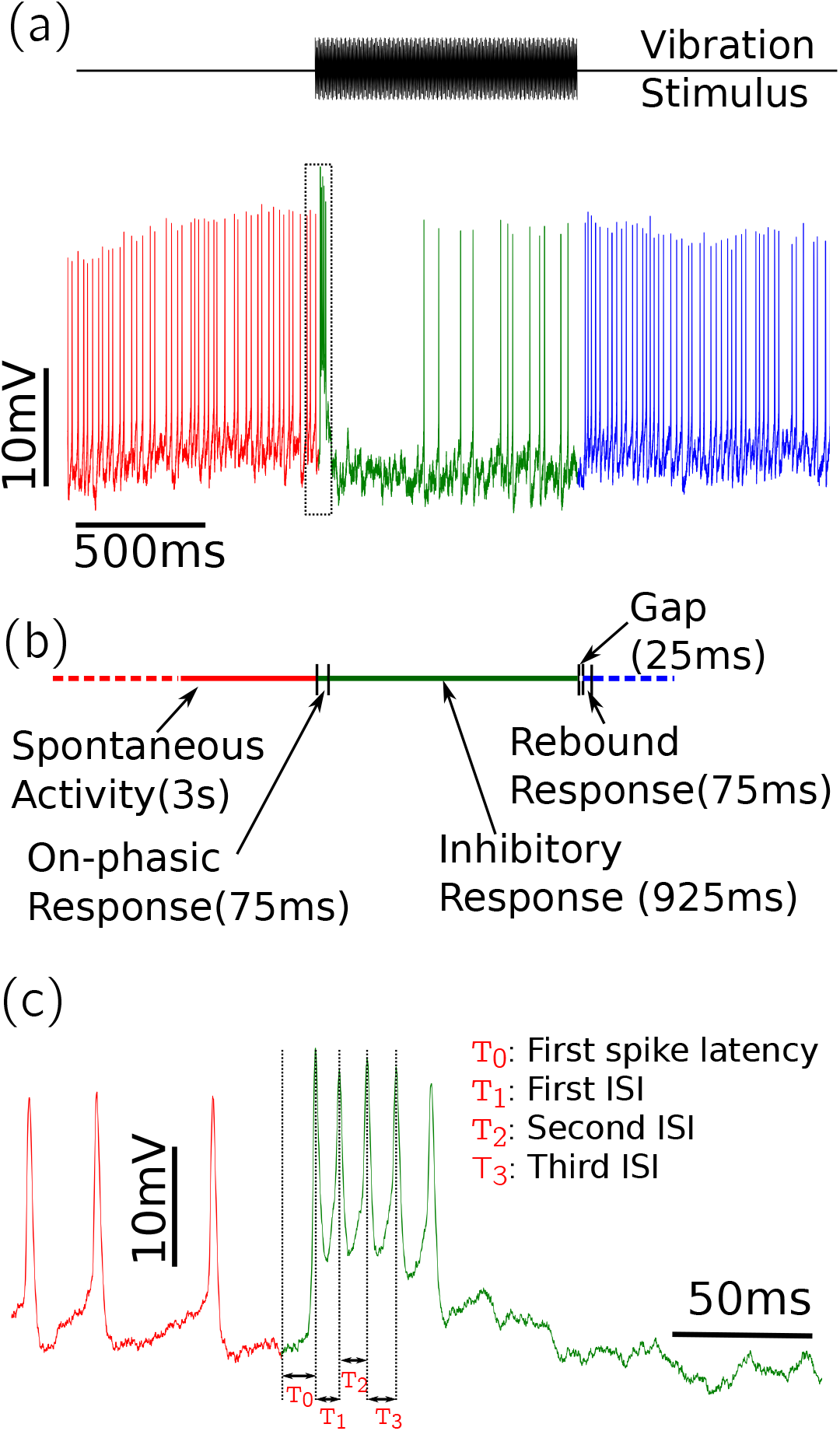
Definition of activity periods and spike timing features of electrophysiological responses. **(a)** An example response of DL-Int-1 to 1 s long vibration stimulus of 265 Hz. Activity before stimulus is colored in red, activity during stimulus in green and activity after stimulus in blue. **(b)** The definitions of the four activity periods used for analyzing electrophysiolgical properties of DL-Int-1. **(c)** The trace contained in the dotted rectangle in (a) is magnified and the four spike timing features, *T*_0_, *T*_1_, *T*_2_ and *T*_3_ are defined on it.

**Figure 5.**
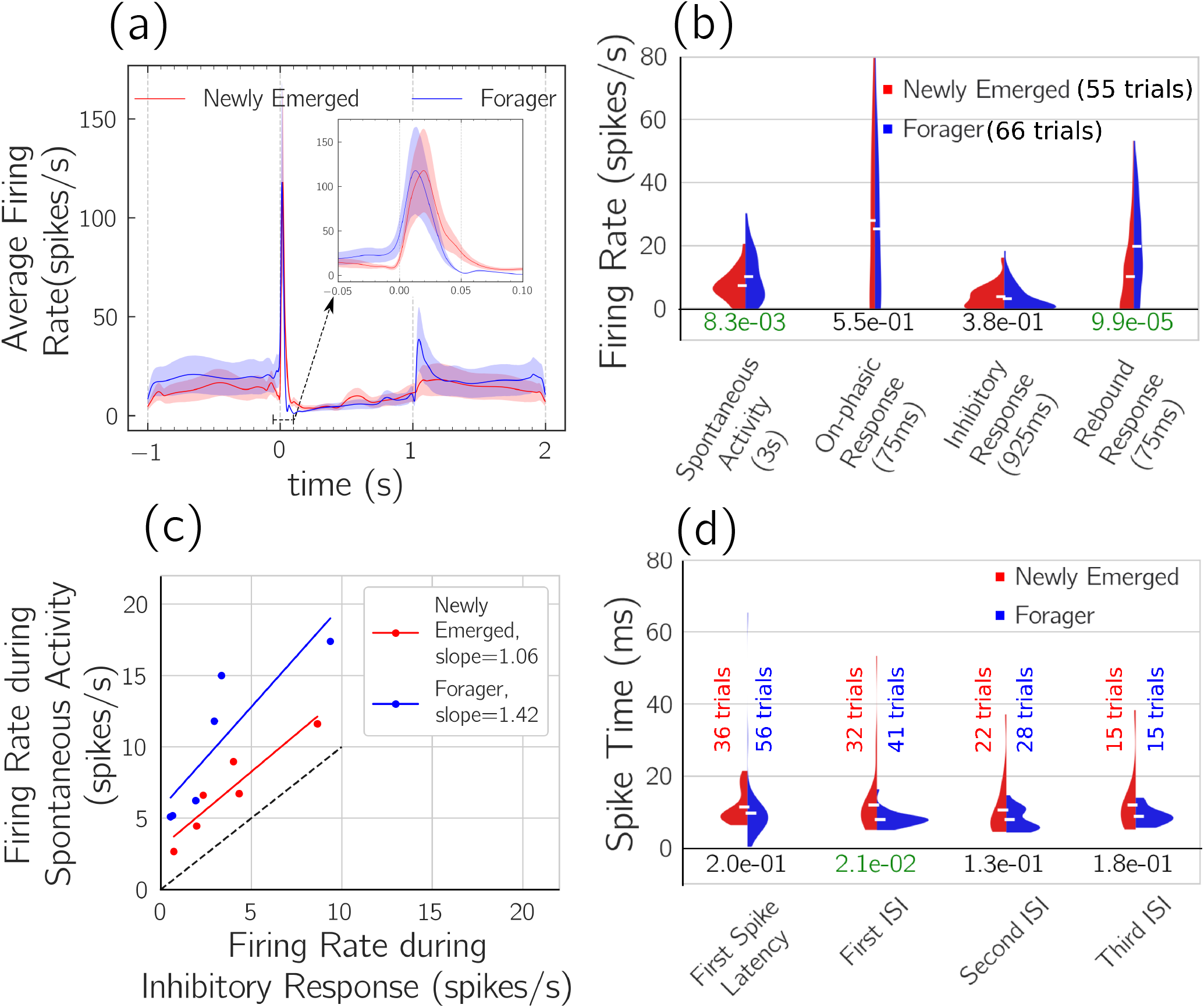
Analysis of electrophysiological properties. **(a)** Comparison of the average firing rate profiles of newly emerged adult and forager DL-Int-1 neurons. Smoothed estimates of time-resolved average firing rates were calculated from responses aligned to stimulus onset using adaptive kernel density estimation (Shimazaki and Shinomoto 2010). Solid lines indicate average firing rates, while shaded regions indicate 95% confidence intervals. Inset: Average firing rate during on-phasic response with expanded time scale. **(b)** Comparison of firing rates during four activity periods. The filled areas represent firing rate distributions estimated using Kernel Density Estimation (see Methods→Analysis of electrophysiology). The distributions were normalized to have equal areas. Horizontal white markers indicate mean values of the distributions. The numbers below the distributions are P-values for the difference of means calculated using Welch’s unequal variance t-test. P-values lower than 5% are highlighted in green. Firing rates during spontaneous activity and rebound response showed significant increases. **(c)** Comparison of the strength of inhibition relative to spontaneous activity by plotting the firing rates during the two periods against each other. Lines were fit using linear least squares regression. The dashed line indicates the line of slope 1. **(d)** Comparison of the timing of the first four spikes of the response using first spike latency, first inter-spike-interval (ISI), second ISI and third ISI. The number of trials that contained enough spikes to calculate the temporal features are shown above the distributions. The distributions were estimated using Kernel Density estimation and normalized to have the same area. The numbers under the distributions are P-values for difference of means calculated using Welch’s unequal variance t-test. The timing of the first four spikes during on-phasic response did not show significant changes, except for the first ISI.

DL-Int-1 is GABAergic and likely part of a disinhibitory network (Ai et al. 2017; Kumaraswamy et al. 2017). Therefore, increased spontaneous rate and post-stimulus rebound in foragers compared to newly emerged adults is expected to result in enhanced strength of the inhibitory signal indicating antennal vibration. To quantify the signal strength of inhibition in DL-Int-1 relative to the level of spontaneous activity, we calculated Relative Inhibition (see Methods→Analysis of electrophysiology) by plotting the firing rates during the two response periods against each other (Figure 5c). In general, higher spontaneous spiking was associated with higher spike rates during inhibition in DL-Int-1 neurons, but the difference in firing rates between spontaneous activity and inhibitory response was larger in foragers than in newly emerged adults. Quantitatively, Relative Inhibition was 0.42±0.24 in newly emerged adults, but 0.66±0.18 in foragers (mean±std error; p-value: 1.34%, Welch’s unequal variance t-test). Additionally, least squares regression in Figure 5c indicated that the difference became larger with firing rate level in foragers (slope of 1.42 for foragers vs 1.06 for newly emerged adult), indicating that foragers had stronger inhibition for a given level of spontaneous activity.

In addition to changes in activity levels during different periods, comparison of firing rate profiles during on-phasic response (Figure 5a inset) indicated a change in the timing of the excitation peak. We investigated this by comparing the first spike latency, first Inter-Spike-Interval (ISI), second ISI and third ISI during on-phasic response (Figure 5d). All four spike timing features showed a systematic reduction during maturation, with an average reduction of 1.76 ms in first spike latency, and 4.01, and 3.09 ms respectively for the first, second and third ISI values. However, none of the changes were highly significant, with first ISI values showing the lowest p-value of about two percent. Thus, spike timing during on-phasic excitation showed a systematic reduction in forager DL-Int-1 neurons, with neither the first spike latency nor the first three inter-spike-intervals showing large significant changes.

## Discussion

In this study, we have compared morphological and physiological properties of an identified vibration-sensitive interneuron, DL-Int-1, between newly emerged adult and forager honeybees. While comparisons of whole cell scalar morphometric measures showed no major differences in broad dendritic structure and gross morphological features, detailed spatial analyses revealed localized changes in the dendritic arborization indicating pruning in the proximal regions and outgrowth in distal regions. The results are consistent with findings of previous studies of changes during maturation in the honeybee antennal lobe (Devaud and Masson 1999) and mushroom body (Farris et al. 2004), which concluded that most of the process of dendritic maturation is completed before emergence, while minor age-dependent and age-independent changes continue for the first few weeks. Such region-dependent changes have also been shown in dendritic arborizations of Kenyon Cells in the adult honeybee (Farris et al. 2001) as well as in the paper wasp (Jones et al. 2009). Comparison of the electrophysiological responses of DL-Int-1 to vibration stimuli between newly emerged adult and forager honeybees showed increased spontaneous activity and stronger post-stimulus rebound, while the qualitative pattern of response remained unchanged. Similar response enhancements have been reported for odor representation in honeybees (Wang et al. 2005), where odor-dependent activity patterns in antennal lobe glomeruli were similar in newly emerged adult and forager honeybees, with older neurons showing higher spiking rates and more active glomeruli.

### Relevance of observed changes

Among scalar measures quantifying dendritic branching and complexity, average partition asymmetry showed a reduction in the DB, while the number of bifurcations did not change significantly. This indicates a redistribution of dendritic branches along with a change in the broad topological structure in the DB. In contrast, for the VB, maximum centrifugal order and Hausdorff dimension reduced significantly without changes in average partition asymmetry. This, together with the large observed changes in dendritic length, surface and volume (Table S4) indicates a reorganization of dendritic branches accompanied by shortening and pruning of dendrites, as well as a reduction in dendritic complexity during maturation of the honeybee from the newly emerged to the forager state. Fine scale spatial analysis of DL-Int-1 using 3D voxels revealed changes in LDL consistent with such morphological adaptations.

The observed changes in neuron morphology of DL-Int-1 are consistent with a refinement process during maturation that leads to improved propagation and processing of vibration signals in foragers compared to newly emerged adults. The DL-dSEG region of the honeybee brain is a center for multi-sensory integration, especially for waggle dance signals (Ai and Hagio 2013; Brockmann and Robinson 2007). DL-Int-1 arborizes with fine terminals and boutons in DL-dSEG (Ai et al. 2017), indicating the presence of important synaptic inputs and outputs in the region (Petralia et al. 2016). An increase of dendritic density in the distal regions of the DB could lead to better connectivity with JO afferents and other neurons arborizing in the DL. The broad structure of the DL-Int-1 did not show major changes during maturation, with the dendritic branches of the DB and VB extending to similar regions in the honeybee brain. Under this condition, reduced dendritic length in proximal regions indicates lower electrical resistance for signals propagating through the DB and VB in DL-dSEG. As suggested by computational studies (Rall and Rinzel 1973; Ferrante et al. 2013), this i.e., the lower resistance results in lower signal attenuation. Similarly, the observed increase in dendritic volume in the MB also is consistent with lower signal attenuation in foragers. These effects could be studied in the future using multi-compartmental neuron simulations, since morphological reconstructions for several newly emerged adult and forager DL-Int-1 neurons are available. However, currently, the lack of experimentally determined membrane conductance parameters poses a major hurdle in performing such simulation studies.

The observed changes in electrophysiological responses of DL-Int-1 could reflect an enhancement of relevant response features for downstream processing of waggle dance vibration signals. Spontaneous firing rates were significantly higher in foragers than in newly emerged adults, while firing rates during inhibitory responses were similar. Thus, the inhibitory response to vibration stimuli was relatively stronger in foragers than in newly emerged adults. Since DL-Int-1 is itself inhibitory and possibly part of a disinhibitory network (Ai et al. 2017; Kumaraswamy et al. 2017), this enhancement results in more effective disinhibition. Furthermore, the strength of postinhibitory rebound doubled during maturation. Since inhibition coupled with postinhibitory rebound has been suggested to play an important role in processing temporal signals in insects (Ai et al. 2018) and specifically in detecting temporal features (Hedwig 2016; Alluri et al. 2016; Naud et al. 2015; Yamada et al. 2018), our results indicate improved detection of information encoded in the temporal features of waggle dance sounds in forager honeybees.

### Physiological changes: neuron or network?

The response of DL-Int-1 neurons to vibration stimuli applied to the antennae is the combined effect of its inputs and its own intrinsic electrophysiological properties (Ai et al. 2017). In this study, significant increases were seen in spontaneous activity and the strength of inhibition relative to spontaneous activity, as well as in the strength of postinhibitory rebound. These changes are likely due to maturation of the electrophysiological properties of DL-Int-1 as well as its connected neuronal network. Specifically, the adaptations in the strength of inhibition relative to spontaneous activity could have a stronger dependence on network factors as DL-Int-1 is inhibitory and is believed to be inhibited in turn as part of a disinhibitory network in the honeybee primary auditory center (Ai et al. 2017; Kumaraswamy et al. 2017). Clarification of the contribution of these two sources would be beneficial for further understanding the role of DL-Int-1 in networks that process waggle dance vibration signals in the primary auditory center of the honeybee.

### Genetically programmed aging or foraging experience?

In this study we have quantified morphological and physiological changes in DL-Int-1 as honeybees mature from newly emerged adult bees (1-3 days old) to forager bees (more than 10 days old). There are two major factors that could cause such changes during maturation— genetically programmed aging and foraging experience. Further studies with age-controlled older honeybees with no foraging experience are required to elucidate the effect of these factors on DL-Int-1 maturation.

### Linking observed changes to behavior

After successful foraging, honeybees return to their hive and perform the waggle dance, during which they produce patterns of air vibration pulses. These pulses are detected by follower bees, from which they learn about the location of food sources (Michelsen et al. 1992; Landgraf 2013). There is evidence that DL-Int-1 plays a role in encoding the distance information from waggle dance airborne vibration patterns (Ai et al. 2017; Kumaraswamy et al. 2017). The observed changes indicate that, as the honeybee matures, there is refinement at structural and functional levels of the neurons and networks involved in processing the waggle dance communication signals that prepare the bees to process those important signals as foragers efficiently.

## Acknowledgments

This research was supported by Grant-in-Aids for Scientific Research from the Ministry of Education, Science, Technology, Sports, and Culture of Japan (Grant No. 22570079, 17K00414 and 18K160345); a grant for Challenging Exploratory Research (Grant 15K14569) from the Strategic International Cooperative Program, Japan Science and Technology Agency (JST); a grant from the German Federal Ministry of Education and Research (Grant 01GQ1116); and a grant from the Central Research Institute of Fukuoka University (Grant 151031). We thank Philipp Rautenberg for contributing to early stages of the project and Hiromu Tanimoto for constructive feedback.

## Supplementary data

**Table S1.** Summary statistics of 19 scalar morphmetric measures applied to the whole arborization (WA) subregion of DL-Int-1 morphologies. Columns two and three contain (mean *±* standard deviation) values for newly emerged adult and forager DL-Int-1. Column four contains the p-values for the difference of means calculated using Welch’s unequal variance t-test. Measures with p-values less than 5% are highlighted in red.

| Measure | Newly emerged | Forager | pVal |
| --- | --- | --- | --- |
| Width (along X) ($\mu m$) | $247 \pm 37.2$ | $270 \pm 19.8$ | 0.2517 |
| Depth (along Z) ($\mu m$) | $233 \pm 28$ | $231 \pm 18$ | 0.8953 |
| Height (along Y) ($\mu m$) | $303 \pm 32.7$ | $327 \pm 18.3$ | 0.1982 |
| Avg. diameter ($\mu m$) | $1.27 \pm 0.149$ | $1.28 \pm 0.0845$ | 0.8763 |
| Total dendritic length ($\times 10^4 \mu m$) | $2.12 \pm 0.765$ | $1.94 \pm 0.53$ | 0.6764 |
| Total dendritic surface ($\times 10^4 \mu m^2$) | $8.67 \pm 4.13$ | $7.74 \pm 2.5$ | 0.6798 |
| Total dendritic volume ($\times 10^4 \mu m^3$) | $3.3 \pm 1.94$ | $2.87 \pm 1.08$ | 0.6773 |
| Total number of bifurcations | $736 \pm 377$ | $562 \pm 215$ | 0.3967 |
| Max. Euclidean distance from root ($\mu m$) | $276 \pm 41.1$ | $263 \pm 22.2$ | 0.563 |
| Max. path length from root ($\mu m$) | $555 \pm 130$ | $541 \pm 56.8$ | 0.8328 |
| Max. centrifugal order | $44.7 \pm 15.2$ | $35.2 \pm 8.33$ | 0.2575 |
| Avg. Burke taper | $-0.272 \pm 0.0866$ | $-0.275 \pm 0.103$ | 0.9702 |
| Avg. contraction | $0.859 \pm 0.0105$ | $0.867 \pm 0.0149$ | 0.3367 |
| Avg. bifurcation angle (local) (degrees) | $122 \pm 2.12$ | $123 \pm 2.69$ | 0.4886 |
| Avg. bifurcation angle (remote) (degrees) | $100 \pm 1.31$ | $99.1 \pm 2.55$ | 0.4055 |
| Avg. partition asymmetry | **$0.616 \pm 0.018$** | **$0.584 \pm 0.0252$** | **0.0447** |
| Avg. parent daughter diameter ratio | $0.985 \pm 0.00652$ | $0.982 \pm 0.00822$ | 0.5223 |
| Avg. sibling diameter ratio | $1.12 \pm 0.00696$ | $1.12 \pm 0.0102$ | 0.7947 |
| Hausdorff fractal dimension | $1.44 \pm 0.0753$ | $1.37 \pm 0.0698$ | 0.1268 |

**Table S2.**
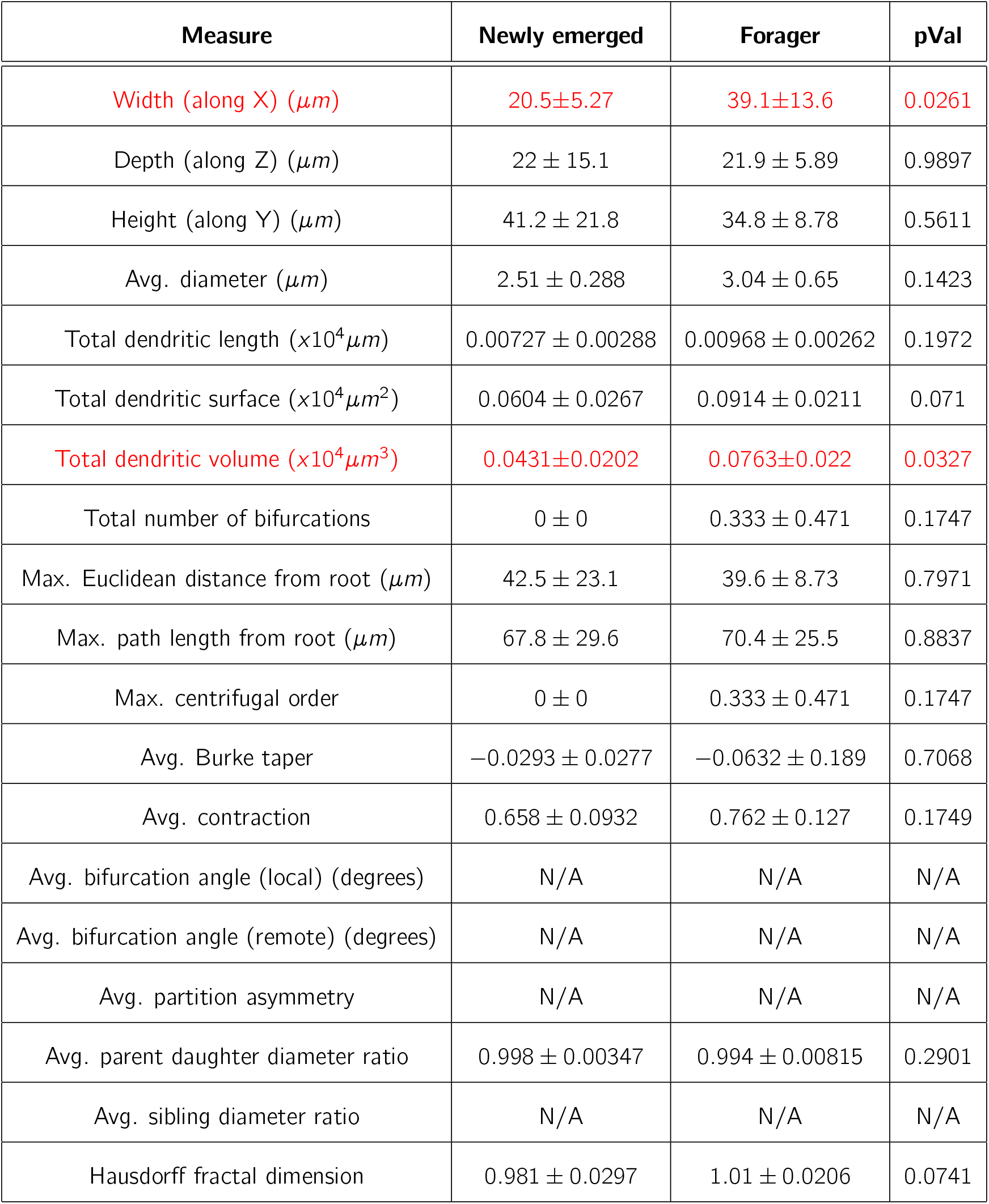
Summary statistics of 19 scalar morphmetric measures applied to the main branch (MB) subregion of DL-Int-1 morphologies. Columns two and three contain (mean *±* standard deviation) values for newly emerged adult and forager DL-Int-1. Column four contains the p-values for the difference of means calculated using Welch’s unequal variance t-test. Measures with p-values less than 5% are highlighted in red. It was not possible to calculate statistics for some measures (marked “N/A”) as one or more morphologies of newly emerged adult or forager DL-Int-1 neurons had no bifurcations in MB.

| Measure | Newly emerged | Forager | pVal |
| --- | --- | --- | --- |
| Width (along X) ($\mu m$) | 20.5±5.27 | 39.1±13.6 | 0.0261 |
| Depth (along Z) ($\mu m$) | 22 ± 15.1 | 21.9 ± 5.89 | 0.9897 |
| Height (along Y) ($\mu m$) | 41.2 ± 21.8 | 34.8 ± 8.78 | 0.5611 |
| Avg. diameter ($\mu m$) | 2.51 ± 0.288 | 3.04 ± 0.65 | 0.1423 |
| Total dendritic length ($\times 10^4 \mu m$) | 0.00727 ± 0.00288 | 0.00968 ± 0.00262 | 0.1972 |
| Total dendritic surface ($\times 10^4 \mu m^2$) | 0.0604 ± 0.0267 | 0.0914 ± 0.0211 | 0.071 |
| Total dendritic volume ($\times 10^4 \mu m^3$) | 0.0431±0.0202 | 0.0763±0.022 | 0.0327 |
| Total number of bifurcations | 0 ± 0 | 0.333 ± 0.471 | 0.1747 |
| Max. Euclidean distance from root ($\mu m$) | 42.5 ± 23.1 | 39.6 ± 8.73 | 0.7971 |
| Max. path length from root ($\mu m$) | 67.8 ± 29.6 | 70.4 ± 25.5 | 0.8837 |
| Max. centrifugal order | 0 ± 0 | 0.333 ± 0.471 | 0.1747 |
| Avg. Burke taper | -0.0293 ± 0.0277 | -0.0632 ± 0.189 | 0.7068 |
| Avg. contraction | 0.658 ± 0.0932 | 0.762 ± 0.127 | 0.1749 |
| Avg. bifurcation angle (local) (degrees) | N/A | N/A | N/A |
| Avg. bifurcation angle (remote) (degrees) | N/A | N/A | N/A |
| Avg. partition asymmetry | N/A | N/A | N/A |
| Avg. parent daughter diameter ratio | 0.998 ± 0.00347 | 0.994 ± 0.00815 | 0.2901 |
| Avg. sibling diameter ratio | N/A | N/A | N/A |
| Hausdorff fractal dimension | 0.981 ± 0.0297 | 1.01 ± 0.0206 | 0.0741 |

**Table S3.** Summary statistics of 19 scalar morphmetric measures applied to the dorsal branch (DB) subregion of DL-Int-1 morphologies. Columns two and three contain (mean *±* standard deviation) values for newly emerged adult and forager DL-Int-1. Column four contains the p-values for the difference of means calculated using Welch’s unequal variance t-test. Measures with p-values less than 5% are highlighted in red.

| Measure | Newly emerged | Forager | pVal |
| --- | --- | --- | --- |
| Width (along X) ($\mu m$) | 215±30 | 259±30.6 | 0.0449 |
| Depth (along Z) ($\mu m$) | 197 ± 40.4 | 224 ± 17.7 | 0.2147 |
| Height (along Y) ($\mu m$) | 221 ± 38.5 | 264 ± 26.2 | 0.0692 |
| Avg. diameter ($\mu m$) | 1.22 ± 0.139 | 1.24 ± 0.071 | 0.7454 |
| Total dendritic length ($\times 10^4 \mu m$) | 1.45 ± 0.587 | 1.55 ± 0.482 | 0.7777 |
| Total dendritic surface ($\times 10^4 \mu m^2$) | 5.65 ± 2.69 | 6.03 ± 2.07 | 0.8105 |
| Total dendritic volume ($\times 10^4 \mu m^3$) | 2.04 ± 1.12 | 2.17 ± 0.821 | 0.8427 |
| Total number of bifurcations | 525 ± 266 | 480 ± 198 | 0.7642 |
| Max. Euclidean distance from root ($\mu m$) | 216 ± 51.2 | 204 ± 11.3 | 0.6232 |
| Max. path length from root ($\mu m$) | 492 ± 124 | 491 ± 80.7 | 0.9885 |
| Max. centrifugal order | 44.7 ± 15.2 | 34.5 ± 8.02 | 0.2255 |
| Avg. Burke taper | -0.306 ± 0.112 | -0.338 ± 0.18 | 0.7439 |
| Avg. contraction | 0.859 ± 0.00858 | 0.867 ± 0.0153 | 0.3419 |
| Avg. bifurcation angle (local) (degrees) | 121 ± 1.69 | 122 ± 2.64 | 0.4063 |
| Avg. bifurcation angle (remote) (degrees) | 102 ± 1.3 | 101 ± 2.57 | 0.4684 |
| Avg. partition asymmetry | 0.619±0.013 | 0.583±0.0306 | 0.0459 |
| Avg. parent daughter diameter ratio | 0.986 ± 0.00574 | 0.983 ± 0.00848 | 0.4515 |
| Avg. sibling diameter ratio | 1.12 ± 0.00595 | 1.12 ± 0.0111 | 0.6279 |
| Hausdorff fractal dimension | 1.37 ± 0.0801 | 1.34 ± 0.0614 | 0.6083 |

**Table S4.** Summary statistics of 19 scalar morphmetric measures applied to the ventral branch (VB) subregion of DL-Int-1 morphologies. Columns two and three contain (mean *±* standard deviation) values for newly emerged adult and forager DL-Int-1. Column four contains the p-value for the difference of means calculated using Welch’s unequal variance t-test. Measures with p-values less than 5% are highlighted in red.

| Measure | Newly emerged | Forager | pVal |
| --- | --- | --- | --- |
| Width (along X) ($\mu m$) | $230 \pm 48.6$ | $240 \pm 23.1$ | 0.6869 |
| Depth (along Z) ($\mu m$) | $184 \pm 47.2$ | $186 \pm 23.8$ | 0.954 |
| Height (along Y) ($\mu m$) | $250 \pm 29.1$ | $265 \pm 36.7$ | 0.5121 |
| Avg. diameter ($\mu m$) | $1.38 \pm 0.189$ | $1.37 \pm 0.153$ | 0.9323 |
| Total dendritic length ($\times 10^4 \mu m$) | $0.656 \pm 0.384$ | $0.373 \pm 0.0848$ | 0.1642 |
| Total dendritic surface ($\times 10^4 \mu m^2$) | $2.96 \pm 1.87$ | $1.62 \pm 0.544$ | 0.1767 |
| Total dendritic volume ($\times 10^4 \mu m^3$) | $1.21 \pm 0.868$ | $0.618 \pm 0.28$ | 0.1938 |
| Total number of bifurcations | $210 \pm 155$ | $81.2 \pm 23.8$ | 0.1218 |
| Max. Euclidean distance from root ($\mu m$) | $236 \pm 21.1$ | $229 \pm 23.9$ | 0.6181 |
| Max. path length from root ($\mu m$) | $458 \pm 118$ | $350 \pm 32$ | 0.0984 |
| Max. centrifugal order | 29.7±10.2 | 15.8±2.34 | 0.028 |
| Avg. Burke taper | $-0.091 \pm 0.0624$ | $-0.0798 \pm 0.0423$ | 0.7468 |
| Avg. contraction | $0.859 \pm 0.0233$ | $0.871 \pm 0.0154$ | 0.3837 |
| Avg. bifurcation angle (local) (degrees) | $123 \pm 3.28$ | $126 \pm 4.31$ | 0.3177 |
| Avg. bifurcation angle (remote) (degrees) | $94 \pm 7.34$ | $90.8 \pm 3.45$ | 0.4141 |
| Avg. partition asymmetry | $0.611 \pm 0.0433$ | $0.573 \pm 0.0468$ | 0.2223 |
| Avg. parent daughter diameter ratio | $0.983 \pm 0.0123$ | $0.979 \pm 0.00892$ | 0.5736 |
| Avg. sibling diameter ratio | $1.13 \pm 0.0201$ | $1.12 \pm 0.0157$ | 0.2153 |
| Hausdorff fractal dimension | 1.24±0.0706 | 1.15±0.0339 | 0.0351 |

**Table S5.**
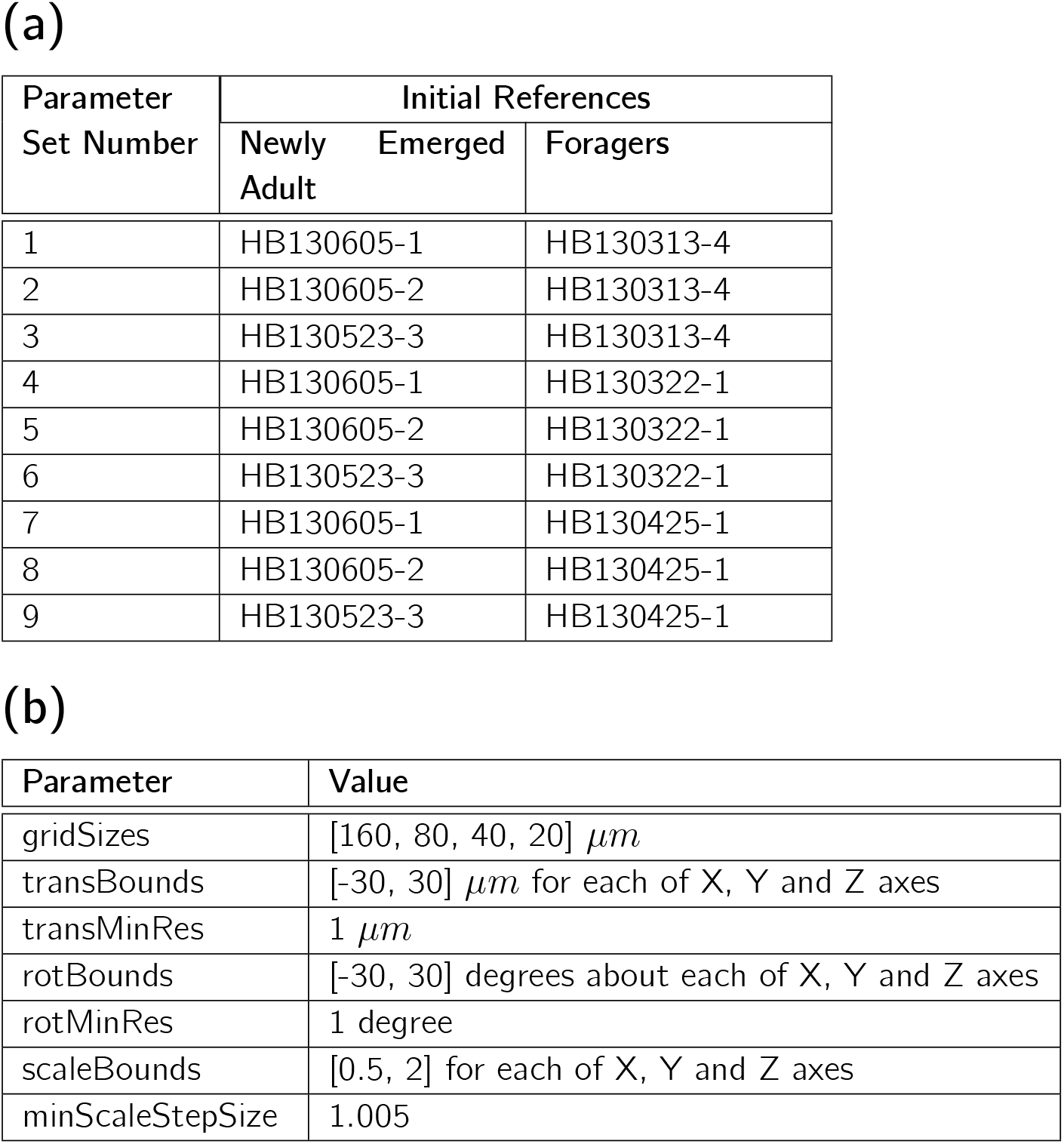
(a) Differing Parameters among the nine parameters sets used for registration. Among the nine parameter sets used for registration, the morphologies used as initial references for co-registering separately newly emerged adults and foragers were different and the experimental identifiers of these morphologies are listed here. Three initial references were used each for newly emerged adults and foragers and taking all possible combinations of these resulted in nine sets of parameters. **(b) Common Parameters among the nine parameters sets used for registration.** Parameters other than the initial references were common among the nine parameter sets. See https://web.gin.g-node.org/ajkumaraswamy/regmaxs/src/master/regmaxsn/core/RegMaxSPars.py for parameter description.

**Figure S1.**
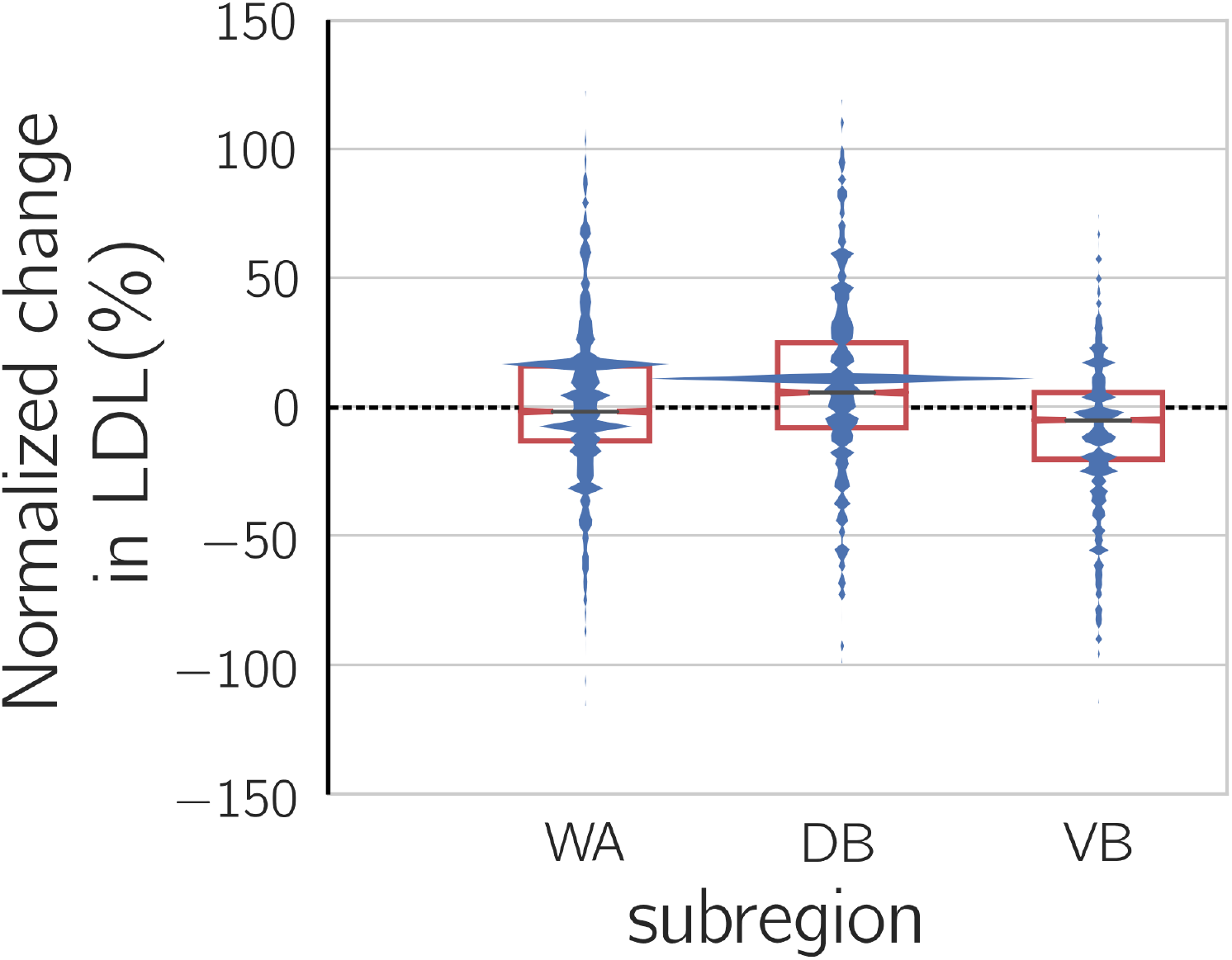
Distributions of normalized change in LDL for WA, DB and VB. The distributions were calculated by smoothing histograms using Gaussian filters with standard deviation equal to 0.001 times the standard deviation of the data. The box plots indicate quartile deviations. The distribution for the DB had more positive values, with values as high as 119%, while the distribution for the VB had more negative values with values as low as −115%. Both distributions had several points lesser than the first quartile and greater than the third quartile.

